# TRPM2 is a Direct Pain Transducer

**DOI:** 10.1101/2025.10.08.681237

**Authors:** Linda Varghese, Mujahid Alizada, Jinquan Yang, Ye Feng, Mitali Malhotra, Xuming Zhang

## Abstract

Chronic pain results from maladaptive interaction between the immune and nervous systems. TRPM2 channels in immune cells (immune TRPM2) are believed to facilitate chronic pain by indirectly promoting immune-inflammatory responses. Whereas TRPM2 in sensory neurons (neuronal TRPM2) acts as a warmth sensor critical to sense innocuous warm temperatures. However, neuronal TRPM2 mediates the warmth sensitivity of less than 3.5% of sensory neurons. The functions of the vast majority (42%) of TRPM2^+^ neurons are unknown. Here we show that neuronal TRPM2 functions as a pain sensor responsible for directly transducing acute and chronic pain independently of immune TRPM2. Both chronic arthritis pain and neuropathic pain were markedly reduced in TRPM2-knockout mice, and the pain deficit was recapitulated by sole deletion of neuronal TRPM2. However, immune and inflammatory responses were largely similar between wild-type and neuronal TRPM2-deficient mice. Moreover, antagonizing joint TRPM2 rapidly reversed chronic arthritis pain without affecting joint inflammation. Mechanistically, TRPM2 is activated by PGE2 and IgG immune complex (IgG-IC) through GαoA and FcγRI coupling, respectively, independently of conventional signalling messengers. Consistently, acute pain induced by PGE2 and IgG-IC was abolished in TRPM2 mutant mice. We conclude that neuronal TRPM2 is a convergent direct pain transducer independently of inflammation, representing an appealing target for alleviating chronic pain.

## INTRODUCTION

Neuroimmune interaction is fundamental to chronic pain(*1*). Activation of either the nociceptive system or immune system mobilises both systems, triggering cooperative responses for removing pathogens and restoring injured tissues. This interacting process often results in chronic pain. For example, nerve injury causes neuropathic pain, while rheumatoid arthritis - an autoimmune disease - produces arthritis pain. The neuroimmune crosstalk suggests common molecular mechanisms underlying chronic arthritis pain and neuropathic pain. Indeed, recent research reported that autoantibodies underlie both rheumatoid arthritis pain and nerve injury-induced neuropathic pain(*2–5*). IgG-IC acts by directly exciting nociceptive neurons through binding FcγRI - a high affinity immune receptor for IgG(*6*) - on sensory DRG neurons, driving arthritis pain and neuropathic pain(*2–5, 7, 8*). However, how IgG-IC excites sensory neurons remains elusive. Furthermore, it is not known whether other common mechanisms co-exist with autoantibodies mediating chronic arthritis pain and neuropathic pain.

TRPM2 is a Ca^2+^-permeable nonselective cation channel mainly expressed in immune cells (immune TRPM2) such as macrophages, monocytes, lymphocytes and neutrophils(*9*). It is activated by Ca^2+^ and the cellular metabolite ADPR functioning as a metabolic sensor(*10–14*). TRPM2 is also a sensor for oxidative stress, though oxidative stress activates TRPM2 indirectly through ADPR produced from the poly ADPR polymerase (PARP-1)/PAR glycohydrolase (PARG) pathway activated during oxidative stress(*15, 16*). Functionally, TRPM2 activation by oxidative stress promotes production of chemokine (CXCL2) and cytokines (IL-1β, IL-6, and TNFα) in monocytes/macrophages aggravating neutrophil infiltration and inflammatory response(*11, 17–20*). It is thus thought that TRPM2 promotes pain indirectly by amplifying immune and inflammatory responses through immune TRPM2(*21, 22*). TRPM2 was also found in sensory DRG neurons (neuronal TRPM2) where it acts a warmth sensor(*23*). However, the warmth-sensitive DRG neurons attributable to TRPM2 are less than 3.5%, despite that around 37% of DRG neurons expressed functional TRPM2(*23, 24*). This raised the question of what the functions of the vast majority of TRPM2^+^ DRG neurons are and whether neuronal TRPM2 acts as a direct pain transducer apart from being a thermo-sensor.

## RESULTS

### Neuronal TRPM2 is crucial to chronic arthritis pain and neuropathic pain

To determine the role of TRPM2 in chronic arthritis pain, we employed antigen-induced arthritis (AIA), an animal model of rheumatoid arthritis. Mice were pre-immunised by methylated BSA (mBSA) followed by intraarticular injection of mBSA into one knee joint(*25*). Joint mechanical pain rapidly developed one day after injection, peaking at around day 3 and then gradually attenuated, but significant joint pain was still evident after 14 days (Fig. 1A). Similar results were obtained with weight bearing tests in which primary joint pain was measured by monitoring body weight distribution difference between the contralateral and ipsilateral limbs of mice (Fig. 1B). Compared to wild-type (WT) mice, joint pain was markedly reduced in TRPM2-knockout (KO) mice (Fig. 1, A and B). We then tested the effect of pharmacological blockade of TRPM2 one day after AIA by injecting the specific TRPM2 antagonist JNJ-28583113 into the knee joint(*26*). Strikingly, joint pain was entirely reversed, and the analgesic effect persisted for two days (Fig. 1, C and D), though no appreciable effect of JNJ-28583113 on joint inflammation was detected (fig. S1A), suggesting that persistent TRPM2 activity in joint sensory nerve endings (i.e. neuronal TRPM2) drives arthritis pain.

**Fig. 1.**
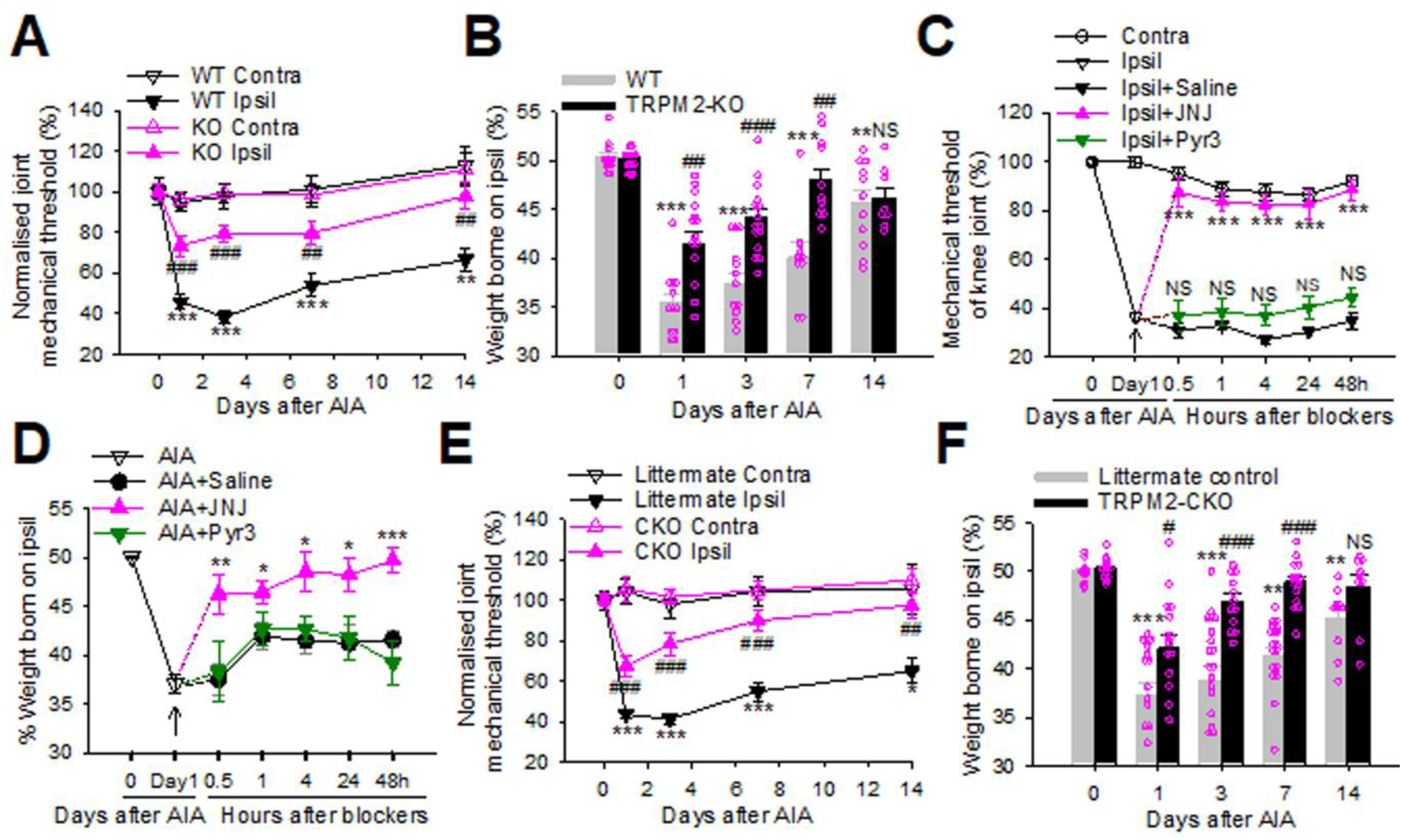
Neuronal TRPM2 channels transduce chronic arthritis pain. **(A)** Mechanical threshold in the contralateral and ipsilateral knee joints in WT and TRPM2-KO mice after AIA induction. n=6-19/group. (**B)** Percentage of body weight borne on the ipsilateral limb in AIA mice. n=8-15 per group. (**C** and **D)** Effects of JNJ-28583113 (2mM), Pyr3 (2mM) and saline on knee joint mechanical threshold (**C**) and body weight distribution (**D**) in AIA mice. n=5-19 per group. arrows indicate drug injection (i.art). ***; NS (not significant), compared to saline. (**E** and **F)** Knee joint mechanical threshold (**E**) and body weight distribution on the ipsilateral limb (**F**) in TRPM2-CKO AIA mice. n=6-19.

To specifically investigate the role of neuronal TRPM2 and exclude the indirect effect of immune TRPM2 on chronic pain, we generated TRPM2 conditional knockout (TRPM2-CKO) mice by crossing homozygous floxed TRPM2 mice (TRPM2^fl/fl^) with Advillin^Cre^ mice in which *Cre* is exclusively expressed in DRG neurons(*27*). We found that TRPM2 protein was expressed in 46.4% (1195/2576) of DRG neurons (fig. S1B), compatible with the expression of functional TRPM2 in 37% of DRG neurons(*23*). 42.9% (513/1195) of TRPM2^+^ DRG neurons co-expressed TRPV1, and 40.8% (498/1219) and 15.9% (118/744) of TRPM2^+^ DRG neurons co-expressed substance P and IB4, respectively (fig. S1, B to E). As expected, TRPM2 was absent in DRG neurons but detectable in joint tissues from TRPM2-CKO mice, whereas TRPM2 was lost in both DRG and joint tissues from TRMP2-KO mice (fig. S1, F to H). Intriguingly, TRPM2-CKO mice reproduced all the pain deficits in TRPM2-KO mice (Fig. 1, E and F), suggesting an essential role for neuronal but not immune TRPM2 in chronic arthritis pain.

We further examined the role of neuronal TRPM2 in neuropathic pain caused by spared nerve injury (SNI), a different type of chronic pain. Mechanical and cold allodynia and heat hyperalgesia were prominent in WT mice 7 days after SNI but all abolished in TRPM2-KO mice (Fig. 2, A to D). All these pain deficits were again recapitulated in TRPM2-CKO mice (Fig. 2, E and H), further consolidating that neuronal but not immune TRPM2 is a predominant player in chronic pain.

**Fig. 2.**
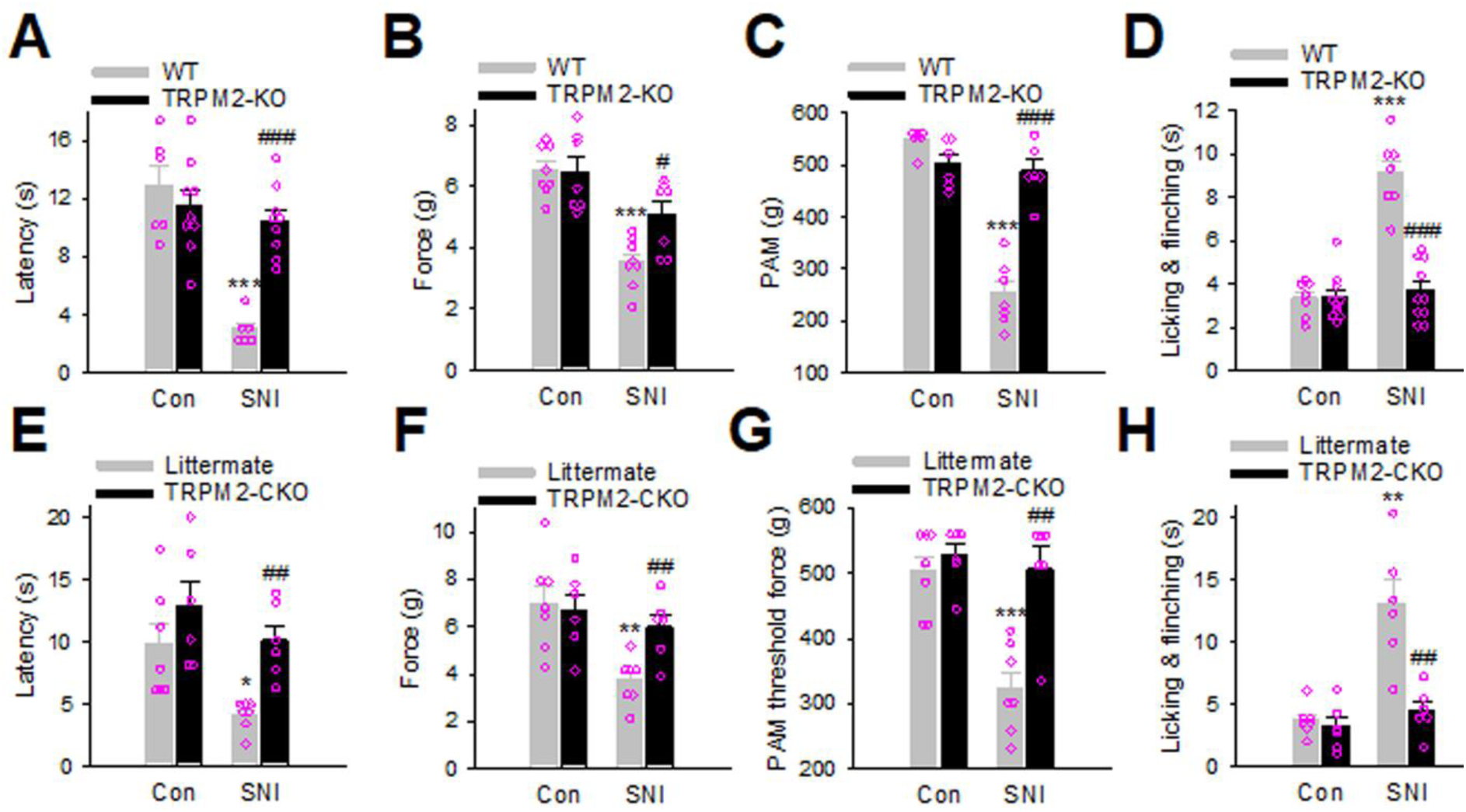
Neuronal TRPM2 channels transduce chronic neuropathic pain. (A to. **H)** Paw withdrawal latency to infrared heat (**A, E**), paw withdrawal force to von Frey filament (**B, F**) and PAM device (**C, G**) and licking and flinching time to acetone (**D, H**) in mice 7 days after SNI surgery in the contralateral (Con) and ipsilateral (SNI) paws in WT and TRPM2-KO mice (**A-D**) and TRPM2-CKO mice and littermates (**E-H**). *, **, *** compared to the contralateral paws/joints; #, ##, ### compared to the ipsil paws/joints in WT or littermate mice.

### Effect of TRPM2 on immune-inflammation and neurogenic inflammation

Immune and inflammatory cells are recognised as important players in chronic arthritis pain(*28*). To determine the effect of TRPM2 on joint inflammation in chronic arthritis, we analysed infiltration of inflammatory cells using specific markers including CD68 for macrophages, Ly6C/G for neutrophils/monocytes, CD3 for lymphocytes and toluidine blue staining of mast cells in the knee joints of mice. We found that joint macrophages were markedly increased, peaking around day 3 (Fig. 3A), consistent with a critical role of macrophages in inflammatory arthritis(*28*). However, deletion of TRPM2 did not prevent macrophage recruitment and even enhanced the recruitment at day one of post-AIA (Fig. 3, A and C). Macrophage recruitment in TRPM2-CKO mice was similar to that in WT mice (Fig. 3, A and C), suggesting that pain deficits in TRPM2 mutant mice are not likely mediated by macrophages. Neutrophils were also robustly increased in the knee joints of WT mice at day one and day 3 but resolved at day 7 of post-AIA (Fig. 3B). Equivalent neutrophil recruitment was also found in TRPM2-KO and -CKO mice and remained prominent at day 7 (Fig. 3, B and D). Similarly, joint lymphocytes were markedly increased in WT and TRPM2-CKO mice but not in TRPM2-KO mice (fig. S2, A and B), implying that immune but not neuronal TRPM2 is crucial to lymphocytes infiltration. We further analysed joint mast cells and DRG macrophages, but no difference was observed between all the mice genotypes (fig. S2, C to F). Furthermore, joint pathology including synovial hyperplasia and cartilage damage was indistinguishable between WT and TRPM2 mutant mice (fig. S3, A to D). Collectively, our data suggest that neuronal TRPM2 is involved in neither joint inflammatory cell infiltration nor joint damage, though immune TRPM2 is critical to lymphocyte infiltration in arthritic joints. Therefore, chronic arthritis pain transduced by neuronal TRPM2 is unlikely mediated by inflammatory cells.

**Fig. 3.**
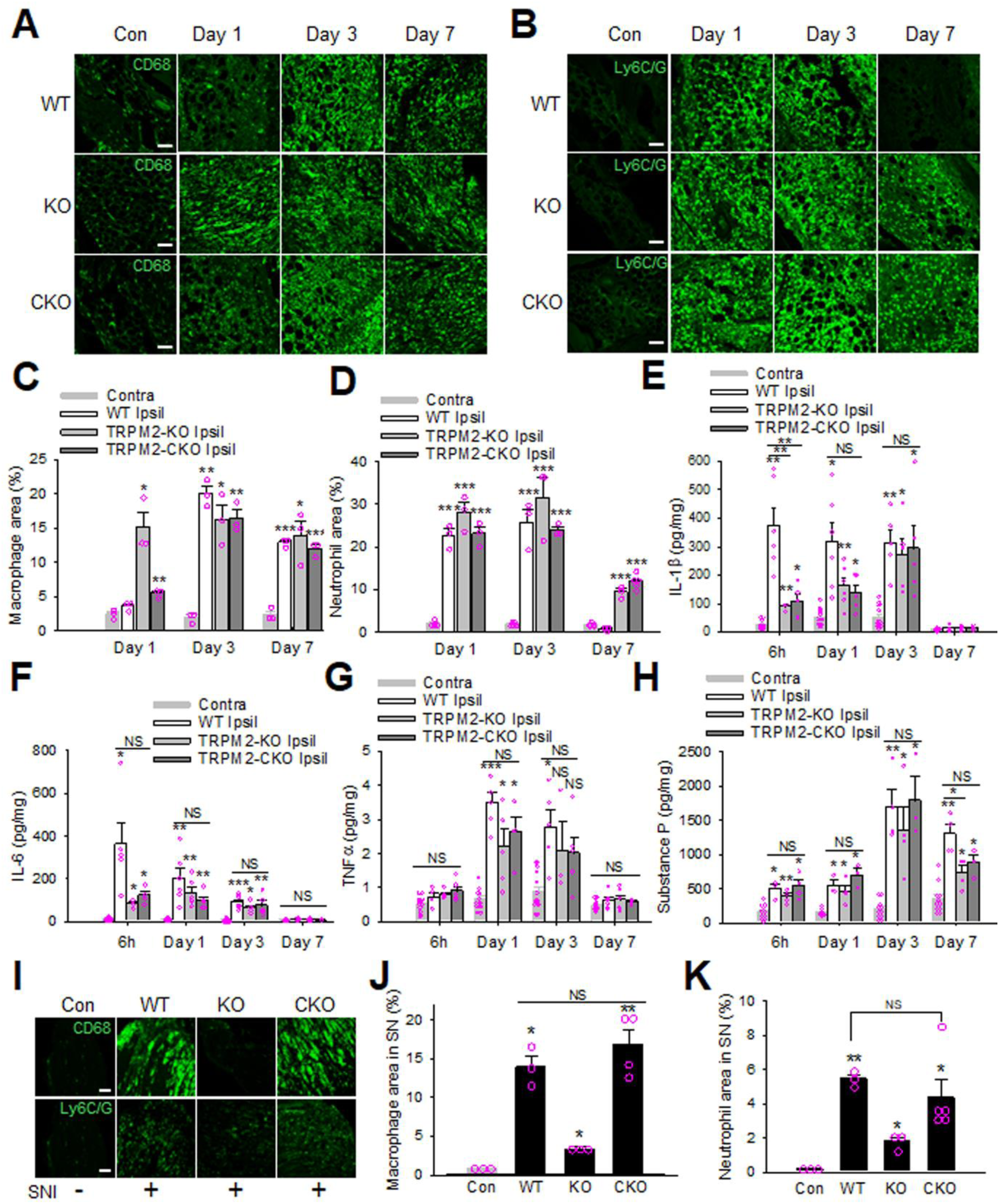
Effect of TRPM2 on tissue inflammation in AIA and SNI models. (**A)** Example images of CD68^+^ macrophages in the synovial tissues of knee joints in different days after AIA. Scale bars, 50μm. (**B)** Representative staining of neutrophils by anti-Ly6C/G in synovial tissues of knee joints from different days of AIA mice. **(C)** Collective results of macrophage-occupied area from similar experiments to those in (**A**). (**D)** Summary of results from experiments similar to those in (**B**). (**E** to **H)** Concentrations of IL-1β (**E**), IL-6 (**F**), TNFα (**G**) and SP (**H**) measured by ELISA in the knee joints of mice after different days of AIA induction. (**I** to **K)** Representative images (**I**) and summary (**J, K**) of CD68^+^ macrophages (**J**) and Ly6C/G^+^ neutrophils (**K**) in the sciatic nerves from SNI mice. Scale bars, 50μm. NS, not significant.

We next analysed major cytokines and neuropeptides critical to joint inflammation and arthritis pain in the knee joints using ELISA assay(*29, 30*). IL-1β and IL-6 were dramatically increased 6h after AIA but TNFα was not increased until one day after AIA in the ipsilateral joints of WT mice (Fig. 3, E to G). However, all these cytokines declined to the basal level 7 days after AIA, suggesting that these cytokines play a role in the early but not chronic stage of arthritis pain. However, no significant difference was found in the production of these cytokines in either TRPM2-KO or -CKO mice except IL-1β at 6h (Fig. 3, E to G). These results suggest that these cytokines are not crucial to TRPM2-mediated arthritis pain, though IL-1β may contribute to the early stage of arthritis pain.

We further examined Substance P (SP) and CGRP, two primary neuropeptides critical to neuroimmune crosstalk implicated in rheumatoid arthrits(*31–33*). SP was markedly increased at all the tested time points, peaking at day 3 but remaining high at day 7 of post-AIA (Fig. 3H), supporting an important role of SP in both acute and chronic stages of arthritis. However, no significant differences were observed between WT, TRPM2-KO and -CKO mice, except that KO mice exhibited a mild reduction at day 7 (Fig. 3H). Surprisingly, there was no change in CGRP between the contralateral and ipsilateral joints of AIA across all mice genotypes (fig. S3E). Thus, TRPM2-mediated arthritis pain is unlikely due to neurogenic inflammation.

To determine the contribution of immune-inflammation to neuropathic pain, similar approaches were used to detect macrophages and neutrophils in the DRG and sciatic nerves 7 days after nerve injury. Consistent with others(*34–37*), DRG macrophages were significantly increased but indistinguishable between WT, TRPM2-KO and -CKO mice (fig. S4, A and B). Macrophages were also dramatically increased at the injury site of sciatic nerves in both WT and TRPM2-CKO mice but absent in TRPM2-KO mice (Fig. 3, I and J). Similarly, neutrophil infiltration was equivalently increased in WT and TRPM2-CKO mice, but not in TRPM2-KO mice (Fig. 3, I and K). These results support that neuronal TRPM2 does not affect the infiltration and recruitment of inflammatory cells at either injury site or DRG during nerve injury but in contrast, immune TRPM2 is indispensable for macrophage and neutrophil infiltration at nerve injury site though it is not required for the recruitment of DRG macrophages. Given that DRG macrophages but not nerve macrophages are critical to the initiation and maintenance of nerve injury-induced neuropathic pain(*34, 37*), our data support that neuronal TRPM2 directly transduces neuropathic pain independent of immune TRPM2 or inflammatory cells.

Taken together, our results demonstrate that neuronal TRPM2 directly transduces both chronic arthritis pain and neuropathic pain with little involvement of immune-inflammation and neurogenic inflammation.

### Neuronal TRPM2 transduces pain signaling evoked by PGE2

We then aimed to understand the common mechanisms of chronic arthritis pain and neuropathic pain carried by neuronal TRPM2. ADPR is an endogenous metabolite agonist for TRPM2 channels and is increased by oxidative stress(*15, 16*). Given that chronic pain is accompanied by oxidative stress, it seems plausible that increased ADPR activates TRPM2 promoting chronic pain. However, neither DRG nor sciatic nerves exhibited increased ADPR in AIA and SNI models (fig. S5, A to D). Therefore, TRPM2-mediated chronic pain is unlikely caused by ADPR.

PGE2 is an important player in the pathogenesis of both rheumatoid arthritis pain and neuropathic pain(*38–41*). Indeed, we found that PGE2 was markedly increased in both DRG and sciatic nerve after nerve injury (Fig. 4, A and B). In AIA model, PGE2 in the knee joints was not altered 6h after AIA but then robustly increased after one day, lasting for over 7 days (Fig. 4C), supporting a key role for PGE2 in sustaining chronic arthritis pain. Comparable increase in joint PGE2 was also observed in TRPM2-KO and -CKO mice and was not significantly different compared to that in WT mice (Fig. 4C). PGE2 was also significantly increased in the ipsilateral DRG one day after AIA induction but then diminished to an insignificant level after 3 days compared to that in the contralateral DRG (fig. S5E). No changes in PGE2 in sciatic nerves were observed in AIA model (fig. S5F). These results suggest that local inflamed joint tissues are a primary source of PGE2 responsible for prolonged arthritis pain and that TRPM2 is not involved in PGE2 production.

**Fig. 4.**
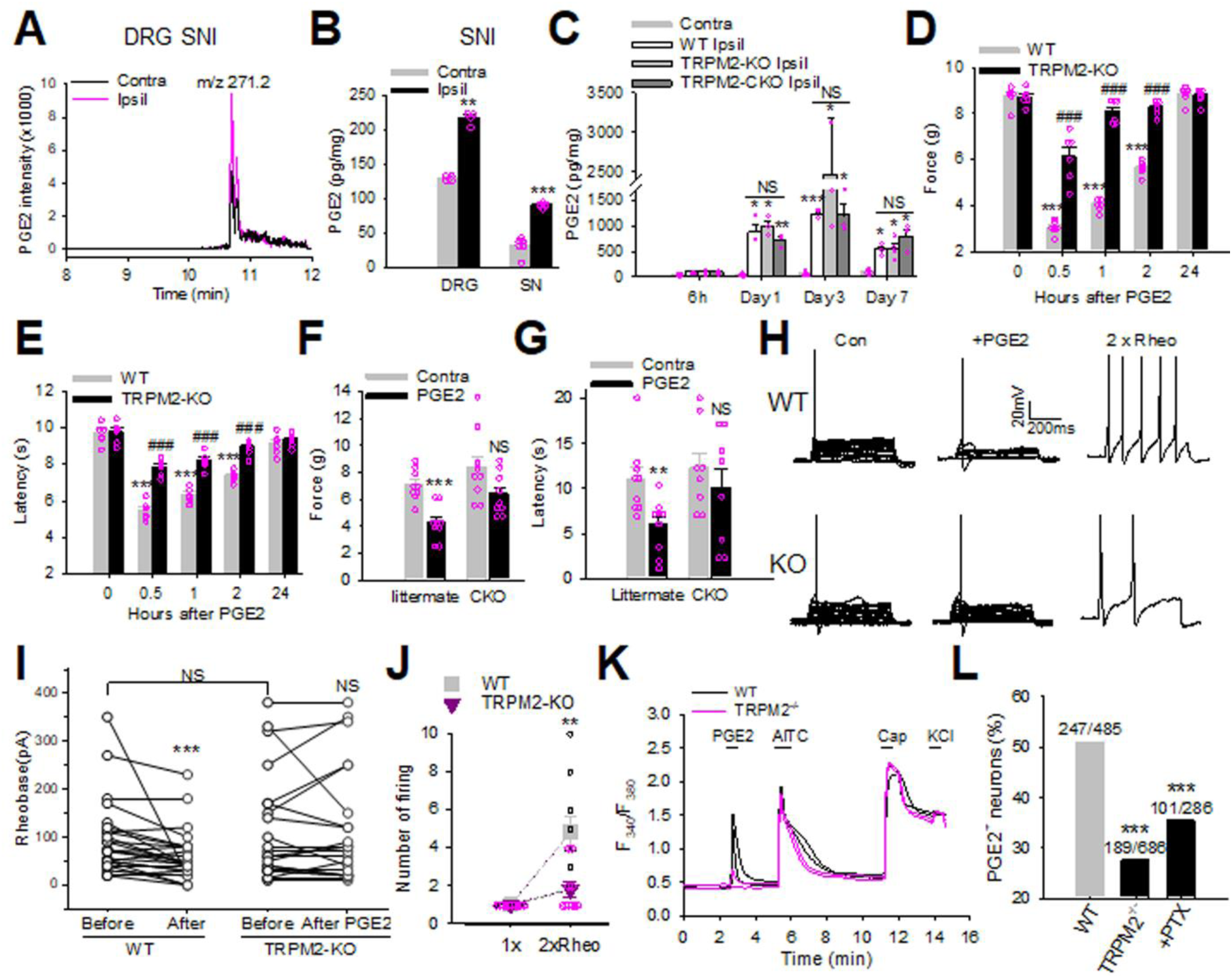
Neuronal TRPM2 mediates PGE2-induced acute pain. **(A)** LC-MS analysis of PGE2 in the contralateral and ipsilateral DRG from SNI mice. (**B)** Summary of PGE2 in DRG and sciatic nerves from experiments similar to those in (**A**). (**C)** PGE2 concentration in the knee joints from mice after different days of AIA induction measured with ELISA. (**D** to **G)** Threshold mechanical force (**D, F**) and withdrawal latency to infrared heat (**E, G**) in the paws of WT and TRPM2-KO (**D, E**), TRPM2-CKO and littmermate mice (**F, G**, 0.5h) after intraplantar injection of PGE2 (10μg/10μl). (**H)** Example firing of DRG neurons from WT and TRPM2-KO mice evoked by depolarization currents steps (left and middle panels) and 2-times rheobase current (right panel) before and after PGE2 treatment (1μM). (**I)** Summary of results from experiments similar to those in (H). (**J)** Summary of number of firing induced by 1x and 2x rheobase current. (**K)** Example Ca^2+^ responses in DRG neurons induced by PGE2 (1μM), AITC (100μM), capsaicin (1μM) and KCl (50mM). **(L**) Summary of PGE2-responding DRG neurons in WT and TRPM2^-/-^ mice.

To determine whether TRPM2 transduces acute pain induced by PGE2, the hind paws of mice were injected with PGE2. Mechanical and heat hypersensitivity were rapidly developed 0.5h after PGE2 injection and then gradually declined to the basal level after 24h (Fig. 4, D and E). However, heightened pain was prevented in TRPM2-KO mice (Fig. 4D and E), and the pain deficit was recapitulated in TRPM2-CKO mice (Fig. 4, F and G), demonstrating that PGE2-induced acute pain is primarily mediated by neuronal TRPM2. At the cellular level, PGE2 enhanced the excitability of sensory DRG neurons, as indicated by reduced rheobase and increased firing frequency caused by PGE2 (Fig. 4, H to J). Neuronal hyperexcitability was, however, abolished in TRPM2-lacking DRG neurons (Fig. 4, H to J). Apart from nociceptor sensitization, PGE2 also triggers spontaneous nociceptor firing, which has been recently shown to be essential to drive mechanical allodynia(*42*). However, neither TRPV1 nor TRPA1 mediate spontaneous firing evoked by PGE2(*42, 43*). We also found that PGE2 induced spontaneous firing in 16.2% (6/37) of WT DRG neurons (fig. S6, A and B). Interestingly, spontaneous firing evoked by PGE2 was rapidly inhibited by blocking TRPM2 with JNJ-28583113, and also significantly reduced in TRPM2-lacking DRG neurons (fig. S6, B to D), suggesting that TRM2 mediates spontaneous firing caused by PGE2. Consistently, the percentage of DRG neurons responding to PGE2 was also markedly reduced in TRPM2-deficient mice (Fig. 4, K and L).

These results support that neuronal TRPM2 directly transduces PGE2-induced acute pain through enhancing the excitability and direct activation of sensory DRG neurons.

### Neuronal TRPM2 transduces acute pain elicited by IgG-IC

Apart from PGE2, recent evidence suggests that IgG-IC is a cause of both arthritis pain and neuropathic pain(*2–5, 7, 8*). Consistently, we found that nerve injury significantly increased IgG immunoreactivity in the ipsilateral lumbar DRG compared to that in the contralateral DRG in WT mice (Fig. 5A). Equivalent increase in IgG immunoreactivity was also evident in TRPM2-KO and CKO mice (Fig. 5B). Similarly, IgG accumulation was drastically increased in the knee joints of mice one day after AIA but then decreased at day 7 (Fig. 5, C and D). Accumulated IgG-IC was not affected by deleting TRPM2 (Fig. 5, C and D). In contrast to the knee joints, IgG immunoreactivity in the DRG was not altered in AIA mice (fig. S7, A and B), suggesting that joint IgG is a primary cause of arthritis pain.

**Fig. 5.**
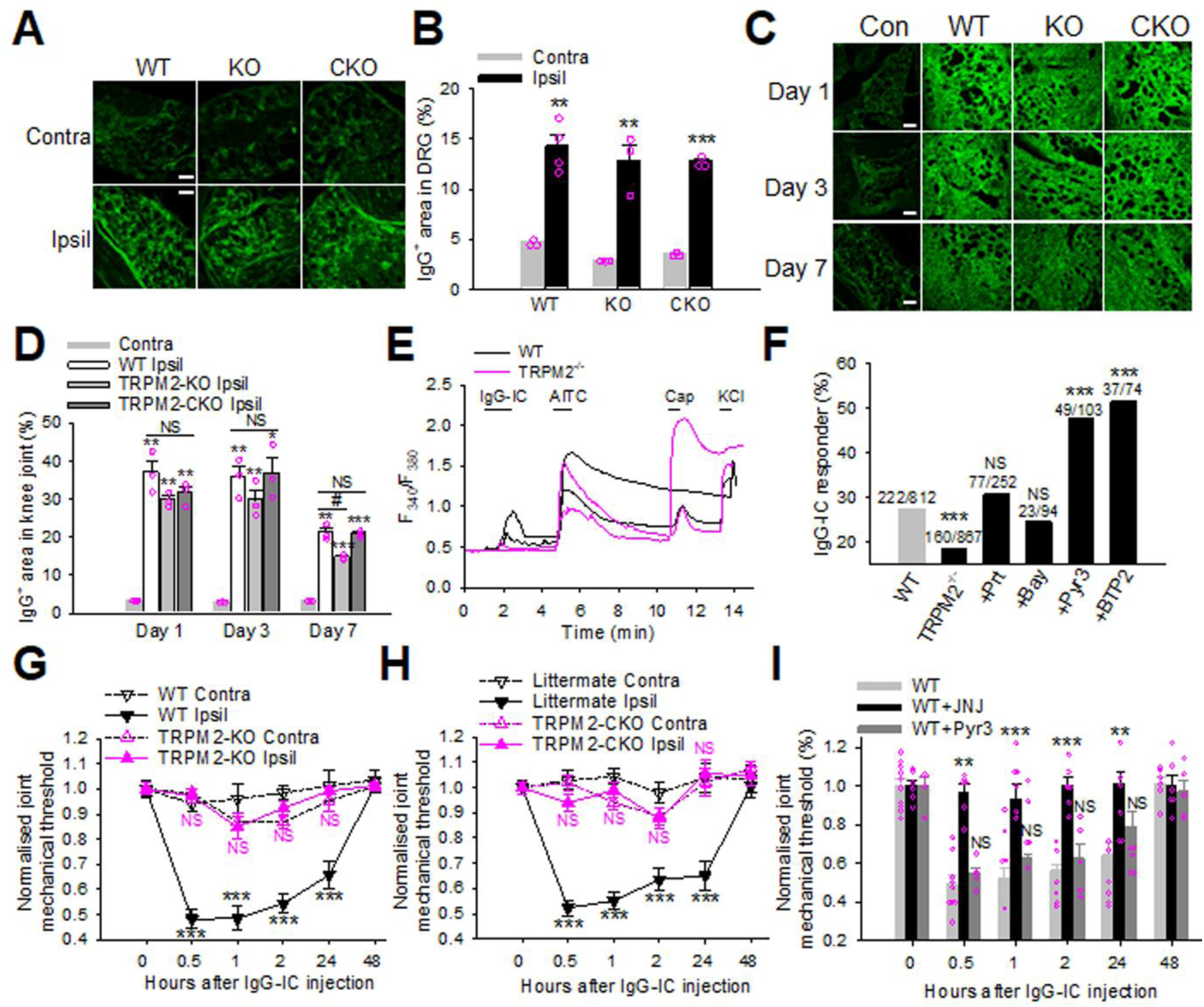
Neuronal TRPM2 transduces acute pain evoked by IgG-IC. **(A)** IgG immunoreactivity in the contralateral and ipsilateral lumbar DRG 7 days after SNI. Scale bars, 50μm. (**B)** Summary of IgG^+^ area in the lumbar DRG. (**C)** Representative staining of IgG immunoreactivity in the knee joint of WT, TRPM2-KO and TRPM2-CKO mice after AIA induction. Scale bars, 50μm. (**D)** Summary of IgG+ staining area from experiments similar to those in (**C**). (**E)** Ca^2+^ responses in WT and TRPM2^-/-^ DRG neurons evoked by IgG-IC (1μg/ml), AITC (100μM), capsaicin (1μM) and KCl (50mM). (**F)** Summary of IgG-IC-sensitive DRG neurons from similar experiments to those in (**E**). **(G** and **H)** Mechanical threshold of the contralateral and ipsilateral knee joints of WT/TRPM2-KO mice (**G**) and TRPM2-CKO mice/littermate (**H**) after intra-articular injection of IgG-IC (2μg). **I** Effect of JNJ-28583113 (2mM) and Pry3 (2mM) on mechanical threshold in the knee joints of WT mice after IgG-IC injection (i.art).

It has been shown that IgG-IC directly excites DRG neurons through activating Syk and TRPC3 channels(*44, 45*). We also found that a brief application of IgG-IC activated 27.3% DRG neurons, 98.7% of which expressed TRPA1 and/or TRPV1 (Fig. 5, E and F). However, we found that IgG-IC-responding neurons were not affected by two different Syk inhibitors Prt062607 and Bay-61-3606 (Fig. 5F). Surprisingly, two specific TRPC3 channel antagonists Pyr3 and BTP2 even enhanced Ca^2+^ responses evoked by IgG-IC (Fig. 5F). Pyr3 and BTP2 also elicited basal Ca^2+^ increases in 41.4% (46/111) and 47.2% (52/110) of DRG neurons, respectively (fig. S8, A and B), which is likely caused by enhanced basal activity of IP3R in the ER due to constitutive inhibition of IP3R by TRPC3(*46*). These results challenge the suggestion of TRPC3 as a downstream channel responsible for IgG-IC-induced neuronal responses(*47*). Nevertheless, IgG-IC responders were significantly reduced in TRPM2-lacking DRG neurons (Fig. 5, E and F), suggesting that neuronal excitation evoked by IgG-IC is mediated by neuronal TRPM2 but not by Syk nor TRPC3.

To determine whether neuronal TRPM2 transduces IgG-IC-induced acute pain *in vivo*, we injected IgG-IC into the knee joints of mice. Joint mechanical pain was rapidly developed 0.5h after injection and lasted for 2 days, though no significant joint inflammation was observed (Fig. 5G and fig. S7C), consistent with the notion that IgG-IC directly excites joint sensory nerve endings triggering joint pain independently of inflammation(*4, 5, 7, 45*). However, joint mechanical pain was all abolished in both TRPM2-KO and TRPM2-CKO mice (Fig. 5, G and H). Moreover, pharmacological blockade of TRPM2 with JNJ-28583113 also entirely reversed acute joint pain induced by IgG-IC, whereas TRPC3 blocker (Pyr3) had no effect (Fig. 5I). Consistently, chronic arthritis pain was not affected by Pyr3 but eliminated by JNJ-28583113 (Fig. 1, C and D). These data strongly demonstrate that neuronal TRPM2 is a predominant channel transducing both acute joint pain and chronic arthritis pain caused by IgG-IC, whereas TRPC3 is dispensable, consistent with a recent report that TRPC3 channels in sensory DRG neurons are not involved in pain transduction(*48*).

Altogether, these experiments demonstrate that neuronal TRPM2 channels directly transduce IgG-IC evoked pain.

### PGE2 and IgG-IC activate TRPM2 channels independently of ADPR

Our behavioural and cellular studies suggest that PGE2 and IgG-IC activate TRPM2 channels mediating acute and chronic pain. To test this idea, we used whole-cell patch clamping to record TRPM2 currents activated by PGE2 in HEK293 cells expressing PGE2 receptors. We found that PGE2 but not vehicle, evoked a large TRPM2-dependent inward current in cells co-expressing EP4 receptor (Fig. 6A). TRPM2 currents were also elicited by activation of EP3 receptors albeit to a lesser degree, but neither EP1 nor EP2 had any effects (Fig. 6B). TRPM2 currents are unlikely caused by Ca^2+^, a known activator of TRPM2 channels(*12, 49–51*), because [Ca^2+^]i was buffered in our recordings (bath solution,1mM Ca^2+^; pipette solution, 2mM EGTA) below 150nM, which is insufficient to activate TPRM2(*50*).

**Fig. 6.**
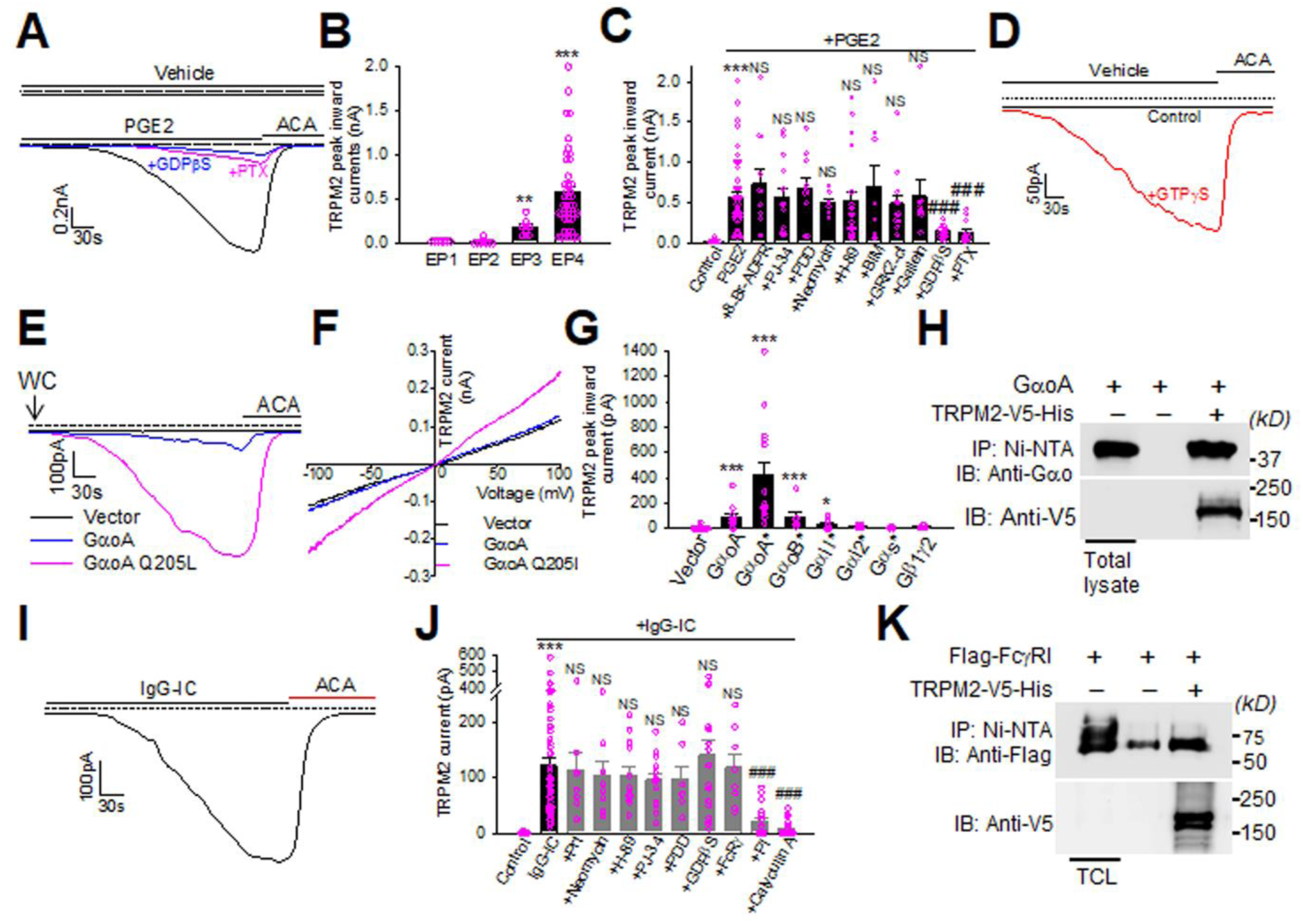
TRPM2 is activated by IgG-IC and PGE2 through noncanonical mechanisms. (**A)** Example current traces in HEK293 cells expressing TRPM2 and EP4 receptor evoked by vehicle and PGE2 (1μM) or after treatment with GDPβS (0.5mM) and pertussis toxin (PTX, 1μg/ml 12h). Current was blocked by *N*-(p-amylcinnamoyl)anthranilic acid (ACA, 20μM). (**B)** Summary of TRPM2 currents mediated by EP1-4 from similar experiments to those in (**A**). (**C)** Collective results from experiments similar to those in (**A**) in cells pretreated with 8-Br-ADPR (300μM), PJ-34 (20μM), PDD00017293 (10μM), neomycin (1mM), H-89 (10μM), BIM (1μM), GRK2-ct, gallein (100μM), GDPβS and PTX. ***, compared to control; NS, ### compared to 2nd bar. (**D)** Example currents recorded from HEK293 cells expressing TRPM2 perfused with vehicle with pipette solution containing GTPγS (100μM) or control solution. € Example currents in HEK293 cells expressing vector, GαoA and GαOA Q205L induced shortly after breaking into whole-cell (WC). Currents blocked by ACA (20μM). (**F)** Average current-voltage curve in HEK293 cells expressing TRPM2 together with vector, GαoA or GαoA Q205L from similar experiments to those in (E). n=8-12. (**G)** Summary of TRPM2 peak inward currents in HEK293 cells expressing TRPM2 together with vector or different G protein subunits similar to those in (**E**). (**H)** TRPM2 binds to GαoA in a Ni-NTA pull down assay. n=3. (**I)** Inward current evoked by IgG-IC (1μg/ml) from HEK293 cells expressing TRPM2 and FcγRI. Currents blocked by ACA (20μM). (**J)** Summary of results from experiments similar to those in (**I)** but treated with Prt062607 (2μM), neomycin (1mM), H-89 (10μM), PJ-34 (20μM), PDD00017293 (10μM), GDPβS (0.5mM), co-expression with FcRγ, phosphatase inhibitor cocktails (1:100) and calyculin A (200nM). ***, compared to control; NS, ###, compared to 2^nd^ bar. (**K)** TRPM2 binds to Flag-FcγRI in a nickel beads pull down assay. n=3.

We then investigated the signalling mechanisms of TRPM2 activation by PGE2. TRPM2 activation was not affected by 8-Br-ADPR, an ADPR antagonist(*52*), nor by PJ-34 and PDD00017273 (Fig. 6C), inhibitors for PARP-1and PARG, respectively, critical to ADPR production. ADPR is thus not involved in TRPM2 activation, consistent with the absent effect of chronic arthritis pain and neuropathic pain on ADPR production (fig. S5, A to D). No effects were observed, either, by inhibiting phospholipase C with neomycin, inhibiting PKA and PKC with H-89 and Bisindolylmaleimide, respectively, nor by inhibiting/chelating Gβγ with gallein and Mas-GRK2-ct (Fig. 6C). However, it was inhibited by inactivating G proteins through dialysing cells with non-hydrolysable GDP analogue GDP-β-S or by inactivating Gαi/o with pertussis toxin (PTX) (Fig. 6C). Conversely, activation of endogenous G proteins by intracellular application of GTPγS was sufficient to activate TRPM2 (Fig. 6D). These results suggest that PGE2 activates TRPM2 through Gαi/o. The conclusion is further supported by our finding that TRPM2 is activated by Gαi/o-coupled EP3 and EP4 receptors but not by EP1 and EP2 receptors coupled to Gαq and Gαs, respectively (Fig. 6B). Collectively, these results demonstrate that Gαi/o signalling is a novel TRPM2 activator independently of Ca^2+^, ADPR and other conventional downstream messengers.

In experiments aiming to identify the Gαi/o subunits responsible for activating TRPM2, we found that GαoA but not vector co-transfection is sufficient to trigger TRPM2-dependent inward currents shortly after breaking cells into whole-cell configuration (Fig. 6E). Constitutively activated GαoA Q205L mutant provoked even more prominent TRPM2 currents and enhanced voltage-dependent gating of TRPM2 (Fig. 6, E and F). Appreciable TRPM2 activation was also triggered by activated GαoB and Gαi1, but other G protein subunits including Gαi2, Gαs and Gβ1γ2 had negligible effect (Fig. 6G). Furthermore, GαoA were found to bind TRPM2 in a nickel beads pull down assay (Fig. 6H), suggesting that GαoA directly activates TRPM2 through physical interaction. In further support of this idea, Gαo proteins were abundantly expressed in DRG neurons(*53*), and inactivating Gαi/o proteins with PTX markedly inhibited Ca^2+^ responses in DRG neurons induced by PGE2 (Fig. 4L). Thus, our results suggest that GαoA is a novel TRPM2 activator mediating neuronal excitation caused by PGE2.

We next examined whether IgG-IC activates TRPM2 through FcγRI, known to mediate chronic arthritis pain and neuropathic pain caused by autoantibodies(*2, 4, 5, 7, 8*). Interestingly, we found that IgG-IC perfusion also activated TRPM2 currents in HEK293 cells expressing FcγRI even in the absence of γ chain (Fig. 6I), essential to transduce FcγRI signalling(*6*), and FcRγ co-expression did not alter TRPM2 activation by IgG-IC (Fig. 6J), suggesting signalling-independent activation of TRPM2 by FcγRI. Consistent with this idea, TRPM2 currents induced by IgG-IC were neither inhibited by Syk2 inhibitor nor by inhibitors for PLC/PKA/PARP-1/PARG nor by inactivating G protein with GDP-β-S (Fig. 6J). These data are also compatible with our finding of absent roles of Syk inhibitors in [Ca^2+^]i responses in DRG neurons induced by IgG-IC (Fig. 5F). Overall, these results suggest that FcγRI activates TRPM2 independent of canonical signalling messengers or G proteins. IgG-IC-induced FcγRI crosslinking may directly activate TRPM2 channels. In support of this idea, TRPM2 bound to FcγRI (Fig. 6K), suggesting that TRPM2 and FcγRI form a receptor-channel complex. FcgRI crosslinking requires receptor dephosphorylation by PP1/PP2A serine/threonine phosphatases(*54, 55*). We found that inhibiting PP1/PP2A with phosphatase inhibitor cocktails and Calyculin A both prevented IgG-IC-induced TRPM2 currents (Fig. 6J), supporting that FcγRI crosslinking is sufficient to activate TRPM2.

Collectively, PGE2 and IgG-IC activate TRPM2 through distinct mechanisms independently of conventional signalling messengers.

## DISCUSSION

Chronic pain is driven by maladaptive neuroimmune responses. As TRPM2 channels are abundantly expressed in immune cells, it has been suggested that TRPM2 contributes to chronic pain by indirectly regulating immune responses through immune TRPM2(*21, 22*). Contrary to this view, we found that TRPM2 in sensory neurons (i.e. neuronal TRPM2) directly transduces acute and chronic pain with little involvement of immune TRPM2. Neuronal TRPM2 thus functions as a direct pain transducer. However, neuronal TRPM2 was also proposed as a sensor for innocuous warm temperatures(*23*). This raised the question of how neuronal TRPM2 transduces both innocuous and noxious signals. Presumably, there exist two distinct subpopulations of TRPM2^+^ DRG neurons transducing pain and temperature, respectively. In support of this possibility, only 3.5% of warmth-sensing DRG neurons are mediated by TRPM2(*23*). The vast majority of remaining TRPM2^+^ DRG neurons may be responsible for pain transduction. More research is required to understand this functional dilemma.

Our research revealed neuronal TRPM2 as a common mechanism transducing two different types of chronic pain: rheumatoid arthritis pain and neuropathic pain. We further uncovered the acting mechanisms of neuronal TRPM2 and found that TRPM2 acts as a convergent pain signalling effector by coupling to FcγRI and EP4, two receptors for autoantibodies and PGE2, respectively, both of which are critical players in rheumatoid arthritis pain, neuropathic pain and many other types of chronic pain(*2, 4, 5, 7, 38, 40, 56*). These two receptor-channel coupling mechanisms may constitute a common output machinery underlying the prominent effect of neuronal TRPM2 in chronic pain.

The discovery of functional coupling TRPM2 to both antibody and PGE2 signalling also resolved two important yet unanswered questions: **First,** autoantibodies generated by immune cells have recently emerged as central mediators of neuroimmune crosstalk and common mechanisms of several different types of chronic pain such as rheumatoid arthritis pain, neuropathic pain, fibromyalgia and low back pain(*2, 5, 7, 8, 56–58*). It has been shown that autoantibodies drive rheumatoid arthritis and neuropathic pain through FcγRI in DRG neurons(*2–5, 8*). But a major unanswered question is how autoantibodies directly excite sensory DRG neurons causing pain. **Secondly,** apart from autoantibodies, PGE2 is also known to be central to inflammatory pain, chronic arthritis pain, neuropathic pain and other pain disorders by sensitizing nociceptive channels, especially TRPV1(*38, 40, 41, 59*). Notably, apart from nociceptor sensitization, PGE2 also induces spontaneous nociceptor firing, which has recently been shown to be essential to drive mechanical allodynia(*42*). However, the mechanisms underlying spontaneous nociceptor firing evoked by PGE2 are unknown. Neither TRPV1 nor TRPA1 channels mediate this process(*42, 43*). Our research demonstrated that activation of TRPM2 channels is a primary channel mechanism mediating the excitatory effect of both antibody and PGE2 signalling; first, acute pain elicited by IgG-IC and PGE2 was abolished by either deleting or blocking neuronal TRPM2; secondly, the same interventions prevented chronic arthritis pain and neuropathic pain; thirdly, Ca^2+^ responses induced by IgG-IC and PGE2 in DRG neurons were reduced in the absence of TRPM2; lastly, IgG-IC and PGE2 are sufficient to open TRPM2 channels through FcγRI and EP4, respectively. Surprisingly, PGE2 and IgG-IC activate TRPM2 independently of ADPR, Ca^2+^ and other classical downstream signalling messengers. Instead, activated GαoA protein by PGE2 and FcγRI crosslinking by IgG-IC were found to be responsible for TRPM2 activation. These noncanonical TRPM2 activation and coupling mechanisms will impact our understanding of the diverse physiology and diseases mediated by TRPM2 channels and autoantibodies beyond the sensory biology and pain fields such as autoimmune diseases, neuroimmune interaction, neuroinflammation, thermoregulation, insulin secretion, stroke, aging and neurodegenerative disease(*60*).

It should be noted that TRPM2 current recordings were performed when [Ca^2+^]i was buffered to prevent the interference of Ca^2+^-dependent activation of TRPM2 (Fig. 6, A, E and I). However, under physiological Ca^2+^ condition, activation of PGE2 and FcγRI receptors may also initiate parallel downstream signalling leading to increased [Ca^2+^]i , which then further boost direct activation of TRPM2 caused by G proteins and FcγRI crosslinking, respectively.

Despite a prominent role of neuronal TRPM2 in transducing chronic arthritis pain, TRPM2 did not affect SP and CGRP release, in contrast to other nociceptive channels such as TRRV1 and TRPA1 critical to neuropeptide release and neurogenic inflammation(*61, 62*). Nevertheless, SP was dramatically increased in AIA mice and persisted longer (>7 days) than other cytokines, suggesting a more important role for SP in the initiation and maintenance of chronic arthritis pain. Consistent with this idea, SP was found to stimulate synoviocyte proliferation and PGE2 release(*31*). PGE2 was also stimulated by IL-1β, IL-6, TNFα and IgG-IC(*63–66*), all of which were critical to chronic pain. Delayed increase in PGE2 in arthritic joints further supports stimulation of PGE2 by these proinflammatory agents (Fig. 4C). Increased PGE2 was then coupled to TRPM2 forming a convergent pain signalling pathway which may underlie the pivotal role of neuronal TRPM2 in pain transduction.

In contrast to SP, CGRP was not increased in the joints of AIA mice. Neither was CGRP altered in the DRG (data not shown), suggesting that CGRP does not contribute to enhanced arthritis pain in AIA model, contrasting with others’ report of involvement of CGRP in rheumatoid arthritis(*32,33*). Different animal models may underlie the difference. Overall, chronic arthritis pain transduced by TRPM2 is unlikely mediated by neuropeptide-mediated neurogenic inflammation.

Role of immune TRPM2 in immune-inflammation has been widely investigated but yielding variable effects. It was initially found that immune TRPM2 expressed in immune cells are pro-inflammatory by promoting release of cytokines and chemokines from monocytes and macrophages exacerbating neutrophil infiltration(*17–19, 67*). However, others found that TRPM2 inhibited cytokines (IL-1β, IL-6 and TNFα) and chemokine (CXCL2) production and inflammatory cell infiltration through inhibiting NADPH oxidase and ROS production, exerting an anti-inflammatory effect(*68–70*). We also found similar opposing effects of TRPM2 depending on tissues. In AIA model, immune TRPM2 promoted recruitment of lymphocytes in the knee joints, but neither macrophage nor neutrophil recruitment in the inflamed joints required TRPM2 (Fig. 3, A to D and fig. S2, A and B). TRPM2 even inhibited macrophage and neutrophil recruitment (Fig. 3, A and B). This inhibitory effect is unlikely mediated by CGRP, because CGRP release was not influenced by TRPM2, even though CGRP was shown to inhibit macrophage and neutrophil recruitment(*71, 72*). Apart from inflammatory cells in the joints, DRG macrophages were reported to be increased in K/BxN serum-transfer rheumatoid arthritis model and proposed as a mechanism of arthritis pain(*32, 73*). However, we did not find difference in macrophage between the contralateral and ipsilateral DRG in AIA model (fig. S2, E and F), consistent with a previous research(*74*). By contrast, DRG macrophage was markedly increased in SNI model (Fig. 3, I and J), supporting that DRG macrophages are a causal mechanism of neuropathic pain but not arthritis pain. However, DRG macrophage recruitment in SNI does not require TRPM2. On the other hand, TRPM2 is indispensable for the recruitment of macrophages and neutrophils in injured sciatic nerves (Fig. 3, I to K). Overall, TRPM2 exerts both pro-(for lymphocytes) and anti-inflammatory effect, which may explain the variable effects of TRPM2 on inflammation.

Different effects of TRPM2 on inflammatory cells seem to be related to their different origins. It has been shown that increased DRG macrophage in SNI and synovial macrophage in rheumatoid arthritis arise from proliferation of tissue resident macrophages, whereas recruited nerve macrophages originate from infiltration of peripheral blood monocytes(*37, 75*), and this process is mediated by immune TRPM2 (Fig. 3J). It thus appears that immune TRPM2 is indispensable for recruitment of blood-derived infiltrating immune cells but not for the expansion of tissue-resident immune cells such as DRG macrophages and synovial resident macrophages. These resident immune cells are highly heterogenous and plastic primarily responsible for producing proinflammatory cytokines in response to ‘on-demand’ signals contributing to chronic arthritis pain and neuropathic pain(*34, 37, 75–77*). However, deletion of neuronal TRPM2 affected neither DRG and nerve macrophages nor synovial macrophages in SNI and AIA models but markedly inhibited chronic neuropathic pain and arthritis pain. We thus concluded that neuronal TRPM2 transduces chronic pain independent of immune TRPM2 and inflammatory cells.

A notable finding in this research is that chronic arthritis pain was completely abolished by pharmacological blockade of TRPM2 channels in the knee joints but partially reduced by genetic deletion of TRPM2. Deletion of TRPM2 may activate unknown compensatory mechanisms leading to incomplete inhibitory effect. Nevertheless, the robust effect of TRPM2 inhibitor on chronic arthritis pain supports that TRPM2 is a highly promising drug target for the treatment of chronic arthritis pain, neuropathic pain and potentially other chronic pain syndromes caused by autoantibodies and PGE2.

## MATERIALS and METHODS

### Study design

This study was designed to investigate the specific role of TRPM2 channels in sensory neurons in chronic pain and the underlying mechanisms. Chronic AIA arthritis pain and SNI neuropathic pain mice models were used to study the effect of loss of TRPM2 in sensory neurons on chronic arthritis pain and neuropathic pain. Their effects on immune inflammatory responses and tissue pathology were examined using ELISA assay and histology. We also analysed the effect of TRPM2 channels on the responses in sensory neurons using calcium imaging and electrophysiology. Electrophysiology was used to determine the role of various agents and signalling messengers in the activation of TRPM2 channels. Sample sizes were determined based on power calculations, previous experiences and publications in our laboratory. Age/sex-matched animals were randomly assigned to different groups, and experimenters were blinded to genotype.

### Animals

All the animal experiments were conducted in accordance with the project license granted by UK home office and approved by the animal welfare and ethical review committee at the University of Warwick. Animals were maintained in IVC cages in a 12h light/dark cycle with humidity and temperature centrally controlled and food and water available *ad libitum*.

TRPM2-knockout mice were obtained from Prof. Peter McNaughton (King’s College London) permitted by Dr. Yasuo Mori (Kyoto University, Japan). TRPM2 floxed (strain no: 029826) and Advillin-Cre (strain no: 032536) mice were obtained from Jackson laboratory. The mice were backcrossed for at least 7 generations in C57/BL6 background. The genotype of transgenic mice was determined with PCR using genomic DNA isolated from ear tissue biopsy samples. Adult mice between the age of 8-16 weeks were used for all the experiments. Both male and female mice were used and assigned to experimental groups by random. The welfare of animals was regularly monitored throughout the experiments.

### Cell culture and transfection

HEK293 cells were cultured in DMEM high glucose (4.5g/L) medium supplemented with 2mM L-glutamine, 100U/ml penicillin, 100μg/ml streptomycin and 10% FBS (ThermoFisher) in a humidified incubator as described previously(*78, 79*). DRG neurons were isolated from adult mice and plated onto coverslips coated with poly-L-lysine (100μg/ml, Merck) as described previously(*78*). The neurons were briefly cultured in DMEM low glucose (1g/L) medium supplemented with and 1% FBS, 2mM L-glutamine, 100U/ml penicillin and 100μg/ml streptomycin and were used for Ca^2+^ imaging and electrophysiology within 8 hours after isolation. Transfection of HEK293 cells with plasmid cDNA was conducted using TurboFect transfection reagent (ThermoFisher) as described previously(*79*). Transfected cells were used for electrophysiology and protein biochemistry analysis within 48h after transfection.

### Molecular Biology

hTRPM2 in pcDNA4/To vector was obtained from Dr. Andrew M. Scharenberg (University of Washigton, USA) and cloned into pCDNA3-V5-His (6x)-TOP vector (ThermoFisher) via NotI and XbaI sites. mTRPM2 in pCDNA3 clone was kindly provided by Prof. Peter McNaughton (King’s College London). hFcγRI in pCMV3 vector with a Flag tag at the C-terminus was purchased from Sino Biological. All the G protein subunits including GαoA, GαoA Q205L, GαoB Q205L, Gαi1 Q204L, Gαi2 Q205L, Gαs Q227L and Gβ1γ2 and PGE2 receptors (EP1-EP4) were purchased from cDNA resource centre. Membrane targeted Mas-GRK2-ct binding to Gβγ devoid of kinase activity was a kind gift of Dr. Stephen R. Ikeda (NIH, USA)(*80*).

### Nickel beads pull down assay

Pull down assay was used to probe interaction of TRPM2 with GαoA or FcγRI and performed as described previously(*78, 79*). Briefly, HEK293 cells co-expressing TRPM2-V5-His and GαoA or FcγRI-Flag were solubilised in lysis buffer containing 20mM Tris-HCl (pH.7.4), 300mM NaCl, 10% glycerol, 40mM imidazole, 0.2mM EDTA plus protease inhibitor cocktail (Merck). Cell lysate was then incubated with Ni-NTA beads (Qiagen) at 4°C overnight. Beads were then thoroughly washed in lysis buffer followed by boiling in sample buffer. Supernatant was then separated in 7.5%SDS-PAGE gel succeeded by protein transfer and detection using anti-GαoA (SC-13532, Santa Cruz) and anti-Flag (Genscript).

### Mouse pain models

Spare nerve injury model was performed as described previously(*81*). Briefly, an incision was made on the skin in the mid-thigh of mice under anaesthesia using isoflurane. A blunt dissection was then made through the bicep femoris muscle to expose sciatic nerve and three distal branches: sural, common peroneal and tibial nerves. Common peroneal and tibial nerves were ligated using Vicryl suture (5–0) followed by transection distal to the ligation. Sural nerve was kept intact. Muscle and skin in the wound were closed with sutures.

Antigen-induced arthritis model was conducted as described by others(*25*). Animals were preimmunised (s.c.) with 200μg methylated bovine serum albumin (mBSA) (Merck) emulsified in 200μl Complete Freund’s adjuvant (CFA) (Merck). Arthritis was then induced by intra-articular injection of 100μg mBSA into the right knee joints one week after immunisation. Arthritis pain was then assessed, and joint tissues were collected at different days after induction of arthritis.

Acute pain model induced by IgG-IC was conducted by intra-articular injection of 2μg IgG-IC in 10μl saline into the right knee joint. IgG-IC was prepared by incubating BSA (10μg/ml) (Merck) with anti-BSA (2mg/ml) (MP biomedicals) in 1:3 ratio in PBS at 37°C for 40min. Acute pain model induced by PGE2 was carried out by intraplantar injection of 1μg PGE2 (Tocris Bioscience) in 10μl saline.

### Behavioural assays

#### Von Frey test

Mechanical pain was measured using dynamic plantar aesthesiometer (Ugo Basile). Animals were acclimatised in a transparent enclosure (9.6 x 9.6 x 14cm) on a metal mesh for at least 30 min before experiments. Mechanical force was applied perpendicularly to the paws of mice using a standardised filament controlled by a touch stimulator. Threshold mechanical force at which a paw withdrawal response is elicited in animals was then automatically recorded by the device. Animals were tested for at least two trials with at least 5min apart between the trials.

#### Hargreaves test

thermal hyperalgesia was monitored using Hargreaves apparatus (Ugo Basile). After full habituation of animals, the hind paw of animals was exposed to an infrared heat source. Once animals feel pain and withdraw their paws, latency to paw withdrawal was automatically recorded.

#### Acetone evaporation test

A drop of acetone (50μl) was directly applied to the hind paws of animals using a syringe. The duration of animals spent on flinching, lifting, licking and biting the paws were recorded.

#### Weight bearing test

The habituated animals were placed on an Incapacitance Tester (Linton Instrumentation) with two hind paws sitting on the two weight sensors, respectively. Body weight borne by each hind limb was automatically recorded and weight distribution between two limbs were calculated.

#### PAM test

primary joint pain was measured using a Pressure Application Measurement (PAM) device (Ugo Basile). A force senor was directly applied to the joints of mice. Threshold force eliciting a limb withdrawal response was recorded by the device.

### Histology

Animals were transcardially fixed by perfusion with PBS followed by 10% formalin solution under anaesthesia with isoflurane. Knee joints were collected and decalcified in 10% EDTA for 30 days followed by treatment with 30% sucrose. The lumbar L4-L5 DRGs, sciatic nerves, and skin samples were also collected and postfixed in 4% paraformaldehyde overnight followed by 30% sucrose overnight for cryopreservation. Tissues were then embedded in OCT compound and sectioned in a cryostat at 12μm. Tissue sections were loaded on poly-lysine coated slides for staining.

### Immunohistology

Tissues sections were blocked and permeabilised in 5% donkey serum and 0.4% Triton X-100 at RT for 30min. They were then incubated at 4°C overnight with the following primary antibodies: TRPM2 (1:200, rabbit, Alomone lab), CD68 (1:100, mouse, 137002, Biolegend), CD3 (1:200, rat, eBioscience), Ly6C/G (1:200, rat,108412, Biolegend), Anti-mouse IgG (H+L) (1:500, goat, Jackson ImmunoResearch), TRPV1 (1:100, mouse, Santa Cruz), Substance P (1: 100, mouse, MAB4375, Bio-Techne), FITC-conjugated IB4 (1:100, L2895, Merck). After wash, sections were next incubated at RT for 2h with fluorescence-conjugated secondary antibodies (1:500, Alexa Fluor 488/594 donkey/chicken anti-rabbit/anti-mouse/anti-rat or Alexa Fluor 488 donkey anti-goat). The stained sections were mounted onto slides for examination using a Zeiss confocal microscopy. A minimum of five images per animal sample and at least 3 animals were obtained for each experimental group. All the images were analysed using Fiji Image J software.

### Haematoxylin & Eosin (H&E) staining

Tissue sections were stained in 0.1% Mayers haematoxylin at RT for 10min followed by washing with tap water. The tissues were then counterstained briefly with 0.01% Eosin solution and sealed by coverslip.

### Toluidine Blue staining

Tissue sections were first briefly treated with 10% neutral buffered formalin solution and then washed in deionised water. They were then stained with 0.02% Toluidine Blue solution for 40 seconds. After dehydration, the tissues were sealed by coverslip using Vectamount solution. Proteoglycan loss, cartilage damage and erosion in the knee joints were quantified and scored based on the “SMASH” recommendations (0–3)(*82*).

### ELISA

Joint tissues were collected from animals and snap frozen immediately in liquid nitrogen. The tissues were then homogenised in 1ml PBS containing Complete EDTA-free protease inhibitor cocktail (Merck) using a pestle and mortar. Protein concentration of tissue extracts was determined using Bradford reagent (Merck). The concentration of IL-1β, IL-6, TNFα, PGE2, substance P and CGRP in joint tissues were next determined using ELISA kits (Bio-techne) in accordance with the manufacturers’ instructions. Briefly, samples and standard in duplicate were added with assay diluent to each well of a microplate followed by incubation at RT or 37°C (for CGRP) for 2-3h. Sample wells were then washed and incubated with mouse IL-1β/IL-6/TNFα/PGE2/substance P conjugates at RT for 2-3h. Substrate solution was then added for colour development at RT for 15-30min. Colour density was finally determined using a microplate reader at 450nm.

### LC-MS/MS

Lumbar DRG (L4-L5) and sciatic nerves were isolated from animals and then homogenised in a lysis buffer containing 100mM Tris-HCl (pH 7.4), 150mM NaCl, 1mM EDTA, 1mM EGTA plus protease inhibitor cocktail (Merck). Samples were then centrifuged at 1000rpm for 5min. The supernatants were then mixed with equal volume of cold methanol on ice for 20min followed by centrifuge at 14000rpm for 10min. The supernatants were then transferred to a Ultrafree centrifuge filter (Merck) and subjected to centrifuge at 11300rpm for 2min. The filtrate was then snap frozen for further analysis of ADPR and PGE2 by LC-MS. Briefly, samples were loaded onto a Waters Acquity HPLC HSS T3 1.8μm 2.1 x 150mm column and then separated with increasing gradient of acetonitrile solution. Eluted molecules were then positively ionised through heated electrospray ionization (H-ESI) followed by separation and detection in a Quantiva triple quadrupole mass spectrometer (Thermo Scientific). ADPR and PGE2 were identified by different retention time, unique mass-to-charge ratio (m/z) and collision energy. The concentrations of ADPR and PGE2 were calculated based on the area under peaks by comparing to a positive standard of ADPR/PGE2 using Skyline software (version 25.1).

### Electrophysiology

Whole-cell patch clamping was performed on DRG neurons and HEK293 cells at room temperature using thin-walled patch pipettes with a resistance between 2-4Ω. Signals were amplified using an Axopatch 200B amplifier and digitised using a Digidata 1440A (Molecular Device). Excitability of DRG neurons was assessed using current patch clamping with pipette solution containing (in mM) 140 KCl, 1 MgCl2, 5.0 EGTA, 2.5 MgATP, 0.5 Li GTP and 10 HEPES (pH 7.4 with KOH) and bath solution containing (in mM): 140 NaCl, 4 KCl, 10 HEPES, 1.8 CaCl2, 1 MgCl2, 5 Glucose (pH 7.4 with NaOH). The action potentials were elicited by injecting current steps with an increment of 10pA in 500ms before and after treatment of DRG neurons with PGE2 (10μM, 4min). TRPM2 currents were recorded using whole-cell voltage patch clamping with pipette solution containing (in mM): 140 KCl, 10 HEPES, 1 MgCl2, 2 EGTA (pH 7.4 with KOH). To determine ADPR and G protein signaling, 8-Br-ADPR (300μM, Biolog) or GDP-β-S or GTP-γ-S (Merck) was included in pipette solution indicated in the relevant figures. Bath solution was similar as above except that 1.8mM CaCl2 was replaced by 1mM CaCl2. Series resistance was compensated (>80%) and cells were held at -70mV. The I-V relationship of TRPM2 channels was examined by delivering 500ms duration of voltage ramps from -100mV to +100mV from a holding potential of 0mV. Data were analysed off-line using Clampfit 11.0 (Molecular device).

### Ratiometric Ca^2+^ imaging

Ca^2+^ imaging was performed as described previously(*78*). Briefly, DRG neurons were loaded with 3.3μM Furo-2-AM (ThermoFisher) at 37°C for 20min. Neurons imaged using a Nikon Ti2 inverted microscope connected to a sCMOS camera (Photometrics) by exposing neurons alternatively to 340nm and 380 LED illuminators (Cairn Research UK). Emission signals were collected at 510nm every two seconds. Cells were continuously perfused with Hanks’ balanced salt solution containing (in mM): 140NaCl, 4 KCl, 10 HEPES, 1.8 CaCl2, 1 MgCl2 and 5 Glucose (pH 7.4). Solution perfusion and drug application were controlled by an automated perfusion system (Warner Instruments). A 10% increase in F340/F380 over baseline was counted as a positive response.

### Statistical analysis

All data are expressed as mean ± SEM. Significance between groups was determined using Student’s *t* test or one way ANOVA when comparisons between multiple groups were required. Two tailored test was used throughout the study. Chi-squared test was performed for Fig. 3i and Fig. 4f. *P*<0.05 was considered to be significantly different. *, #, *P*<0.01; **, ##, *P*<0.01; ***, ###, **P**<0.001.

## Acknowledgements

We are thankful to Dr. Yasuo Mori (Kyoto University) and Dr. Peter McNaughton (King’s College London) for providing TRPM2-KO mice. This study was funded by the UK Medical Research Council (MR/V04077/2), BBSRC (BB/T01668X/2) and Versus Arthritis UK (21971). The research was also supported by a Warwick Ph.D studentship (L.V.) and China Scholarship Council (CSC)-UoW Ph.D studentship (J.Y.). All lab members provided comments on the manuscript.

## Author contributions

L.V. performed behavioural experiments. M.A. J.Y. and M.M. conducted histology. M.A. performed ELISA assay and mass spectrometry. Y.F. and X.Z conducted Ca^2+^ imaging. X.Z. performed electrophysiology. X.Z. conceptualised the research, obtained funding, supervised research, analysed data and wrote the manuscript.

### Data and material availability

All the data associated with this study are included in the paper or the supplementary materials.

### Supplementary Materials

Figs. S1 to S8

