## Supplementary data for "TRPM2 is a Direct Pain Transducer"

### Supplementary figures

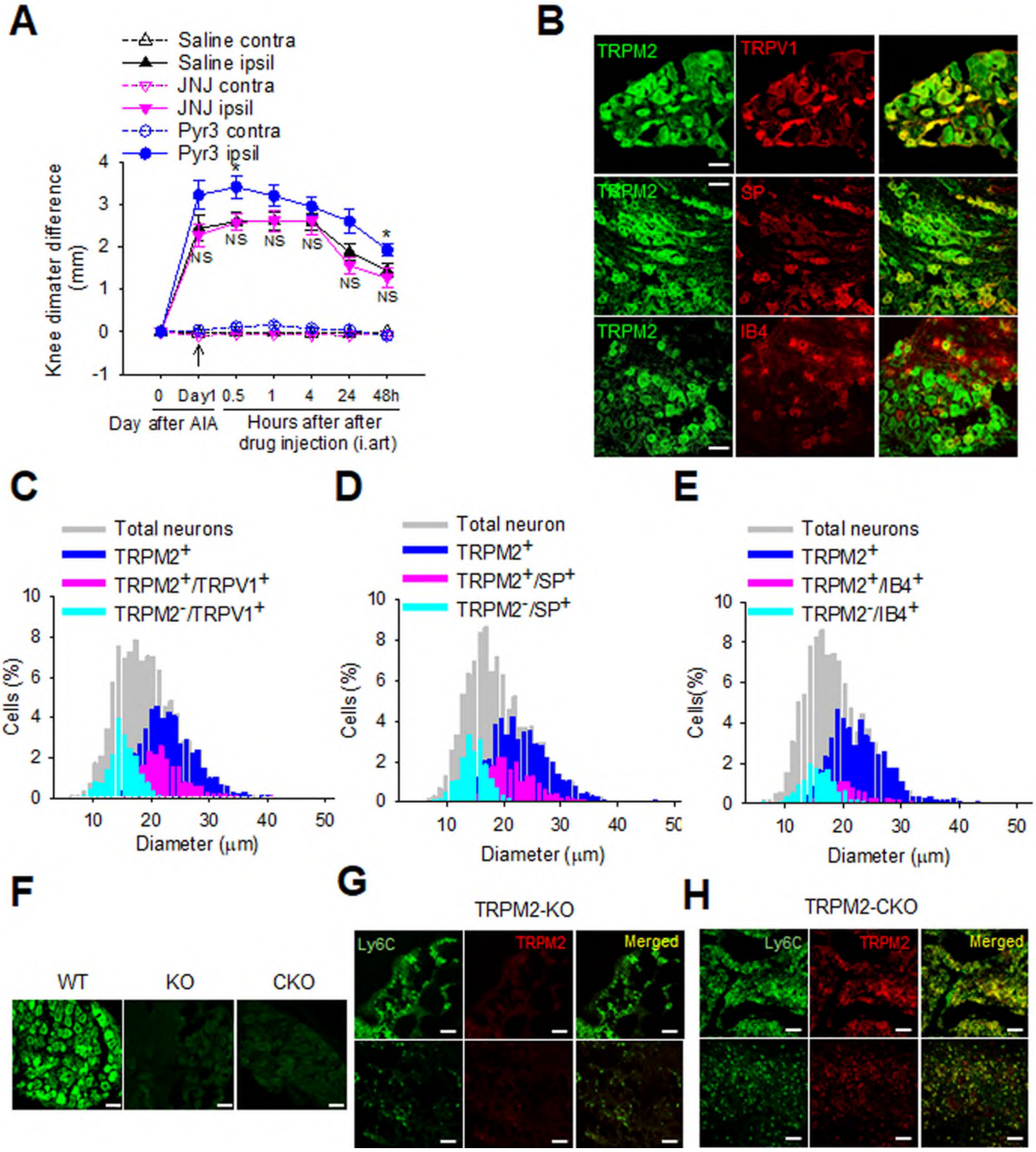


**Fig. S1: TRPM2 is deleted in sensory DRG neurons.** **(A)** Joint diameter difference in mice after AIA induction followed by intraarticular injection of saline, JNJ-28583113 (2mM) and Pyr3 (2mM). Arrow denotes drug administration (i.art). n=5-8 per group. (**B)** Example of immunostaining of TRPM2 with TRPV1(top panel), SP (middle panel) and IB4 (bottom panel) in lumbar DRG neurons. Scale bars, 50μm. (**C-E)** Histogram distribution of $TRPM2^{+}$ DRG neurons with TRPV1-, SP- and IB4-expressing DRG neurons from experiments similar to those in (**B**). $n_{cel}$l=2576 (**C**), $n_{cell}$=2628 (**D**), $n_{cell}$=1605 (**E**). (**F)** Immunohistochemistry staining of TRPM2 in lumbar DRG from WT, TRMP2-KO and TRPM2-CKO mice. Scale bars, 50μm. (**G and H)** Co-expression of TRPM2 (red) with neutrophils/monocytes marker Ly6C (green) in the bone marrow (top panels) and hyper-proliferated synovium (bottom panels) of the knee joints from AIA mice in TRPM2-KO (**G**) and TRPM2-CKO mice (**H**). TRPM2 is deleted in both bone marrow polymorphonuclear cells and synovial inflammatory cells in TRPM2 -KO mice but not in TRPM2-CKO mice. Scale bars, 50μm.

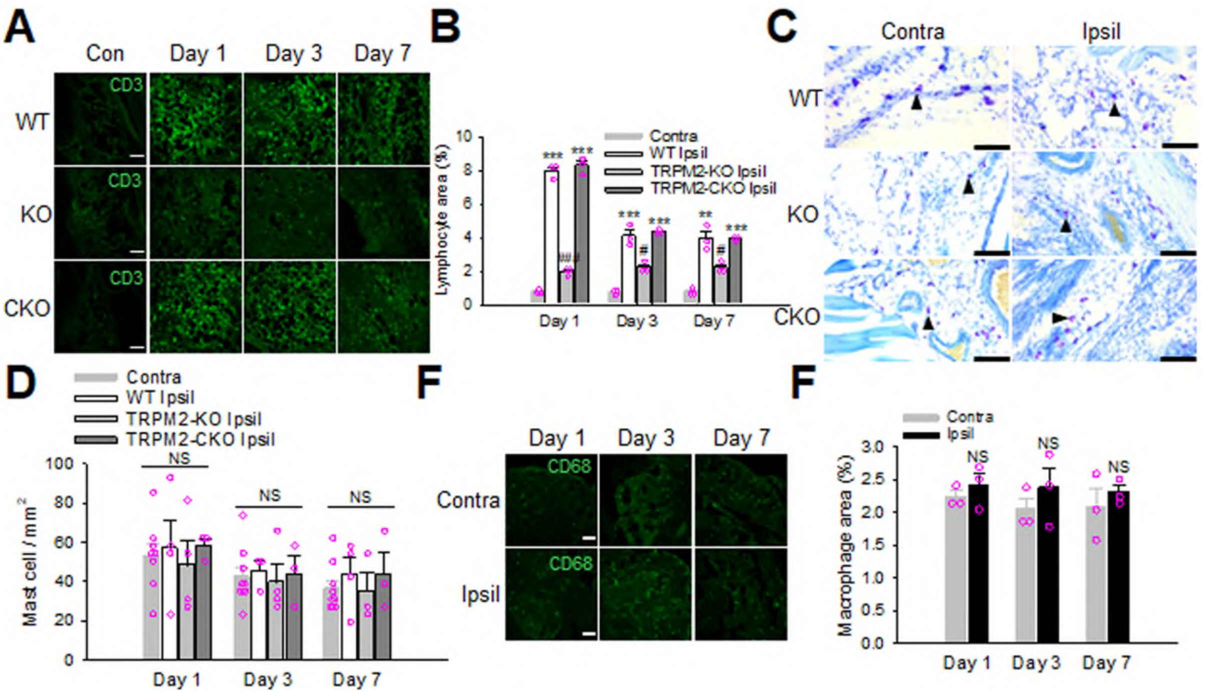


**Fig. S2: Neuronal TRPM2 is not involved in inflammatory cell recruitment in AIA model.** (**A**) Example fluorescence images of lymphocytes stained by anti-CD3 in the synovial tissues of knee joints from WT, TRPM2-KO and -CKO mice at different days after AIA induction. Scale bars, 50µm. (**B)** Summary of percentage of lymphocyte-populated synovial areas from similar experiments to those in (**A**). *** compared to contra; #, ### compared to WT Ipsil. (**C)** Representative images of mast cells revealed by Toluidine blue staining in the contralateral and ipsilateral synovial tissues of knee joints from WT, TRPM2-KO and -CKO mice one day after AIA induction. Arrows indicate mast cells. Scale bars, 100µm. (**D)** Summary of quantification of mast cell density from similar experiments to those in (**C**). (**E)** DRG macrophages stained by anti-CD68 in the contralateral and ipsilateral DRG from mice after different days of AIA induction. Scale bars, 50µm. (**F**) Collective results of macrophage occupied synovial areas from similar experiments to those in (**E**). NS, not significant.

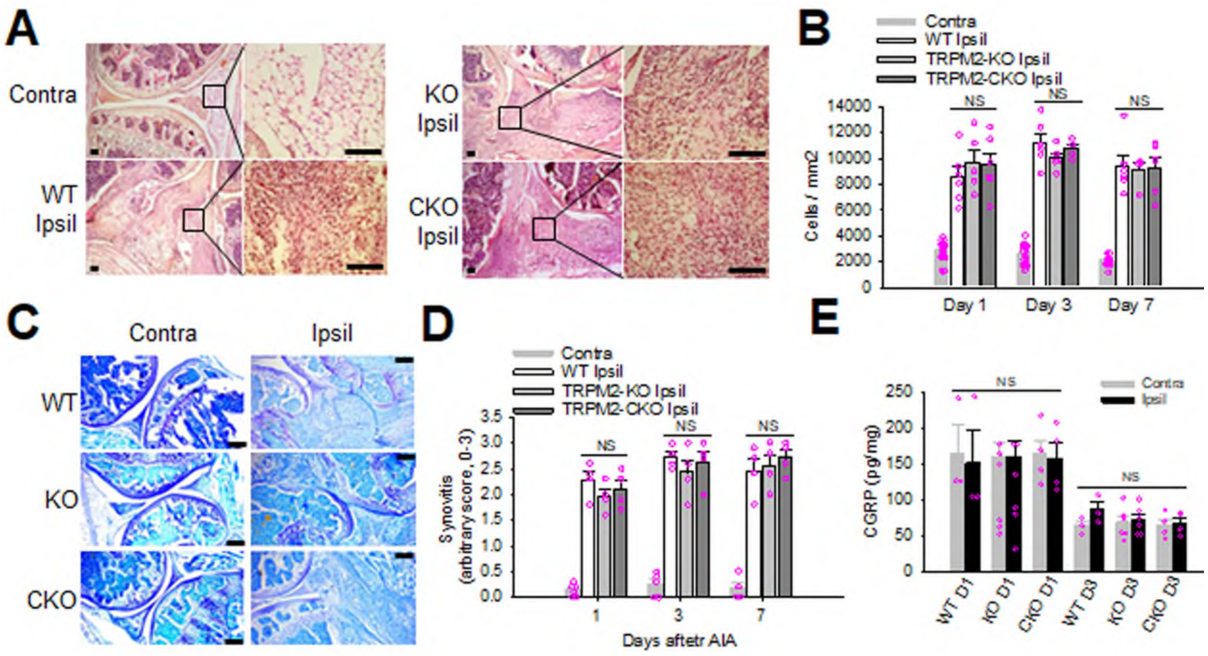


**Fig. S3: TRPM2 has no effect on joint pathology in AIA mice.** **(A)** Representative images of HE staining of the contralateral and ipsilateral knee joints from WT, TRPM2-KO and -CKO mice 3 days after AIA induction. Zoomed synovial tissues indicated by the squared box in the left image panels are shown on the right. Scale bars, 100μm. **(B)** Summary of quantification of cell density in the synovial tissues from similar experiments to those in (**A**). Deletion of TRPM2 does not affect synovial hyperplasia. NS, not significant. **(C)** Toluidine blue staining of the contralateral and ipsilateral knee joints from WT, TRPM2-KO and -CKO mice 7 days after AIA induction. Scale bars, 200μm. (**D)** Summary of scoring of cartilage damage and proteoglycan loss from similar experiments to those in (**C**). TRPM2 deletion does not affect joint pathology. NS, not significant. **(E)** CGRP concentrations in the contralateral and ipsilateral knee joints in AIA mice measured by ELISA.

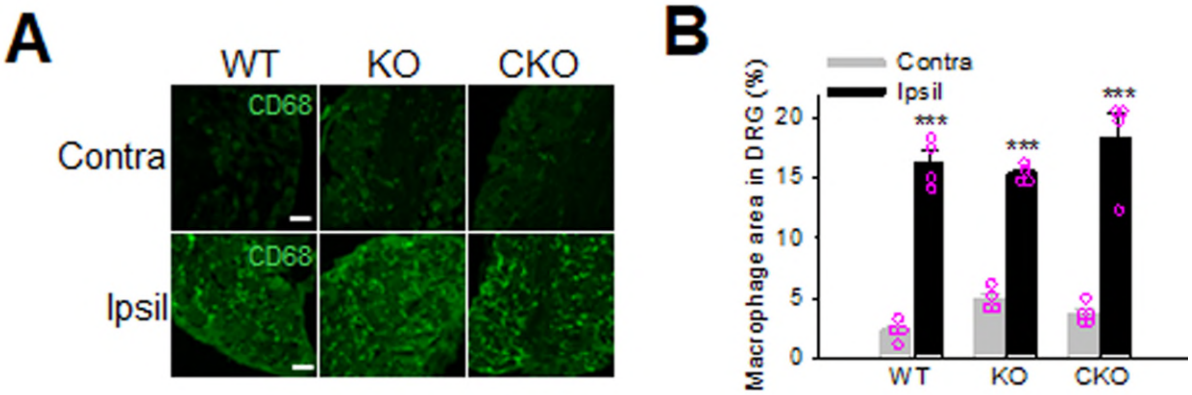


**Fig. S4: TRPM2 does not affect DRG macrophage recruitment in SNI model. (A** and **B)** Example images of DRG macrophages stained by anti-CD68 (**A**) and summary (**B**) of macrophage-occupied areas in the contralateral and ipsilateral lumbar DRG from SNI mice 7 days after nerve injury. Scale bars, 50µm.

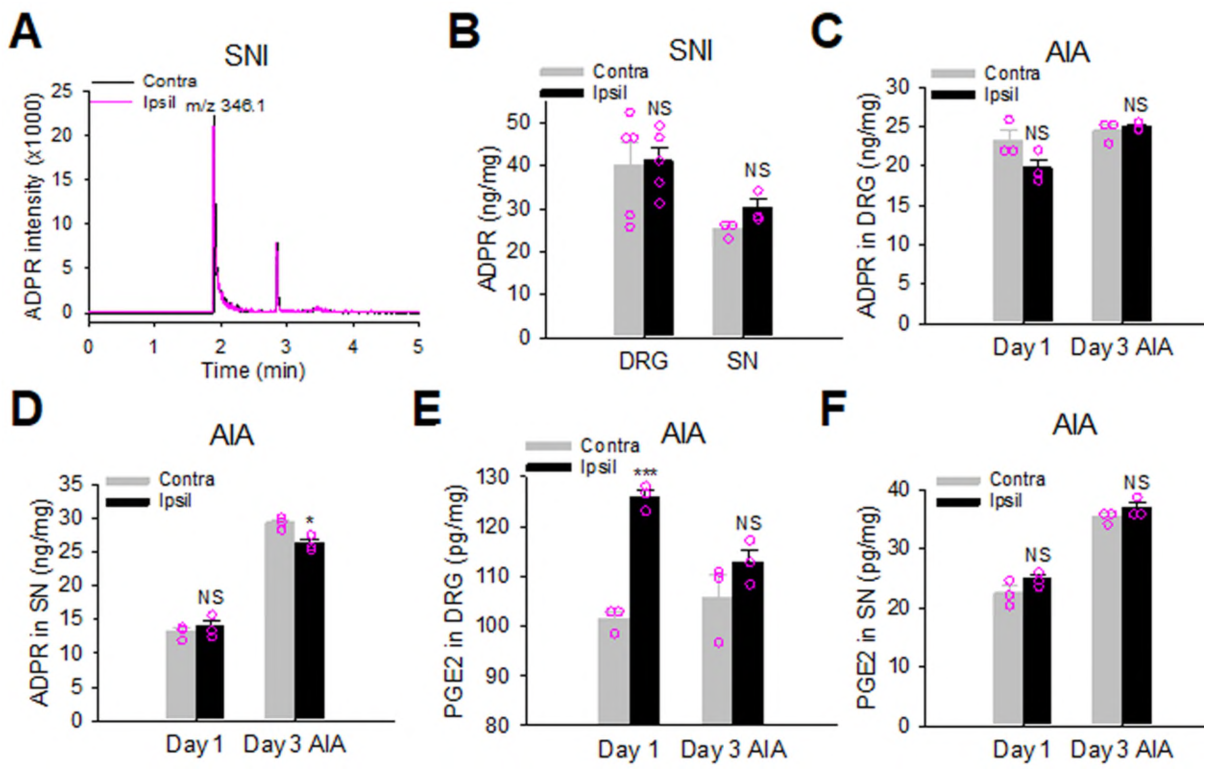


**Fig. S5: ADPR and PGE2 concentrations in the DRG and sciatic nerves (SN) of mice. (A)** Example ADPR peaks from the contralateral and ipsilateral DRG tissues from SNI mice detected by LC-MS. The peaks appear close to 2min after collision at an expected m/z ration of 346.1. (**B)** Summary of ADPR concentrations in the DRG and sciatic nerves (SN) from SNI mice seven days after nerve injury from similar experiments to those in (A). (**C** and **D)** ADPR concentrations in the contralateral and ipsilateral DRG (**C**) and sciatic nerves (SN) (**D**) from mice at Day 1 and Day 3 of post-AIA induction. NS, not significant. *$P<0.05$. ADPR is even reduced in SN three days after AIA induction. (**E** and **F)** PGE2 concentration in the DRG (**E**) and SNs (**F**) from mice after different days of AIA induction measured using LC-MS similar to those in Fig. 3A. *** compared to the contralateral samples. NS, not significant.

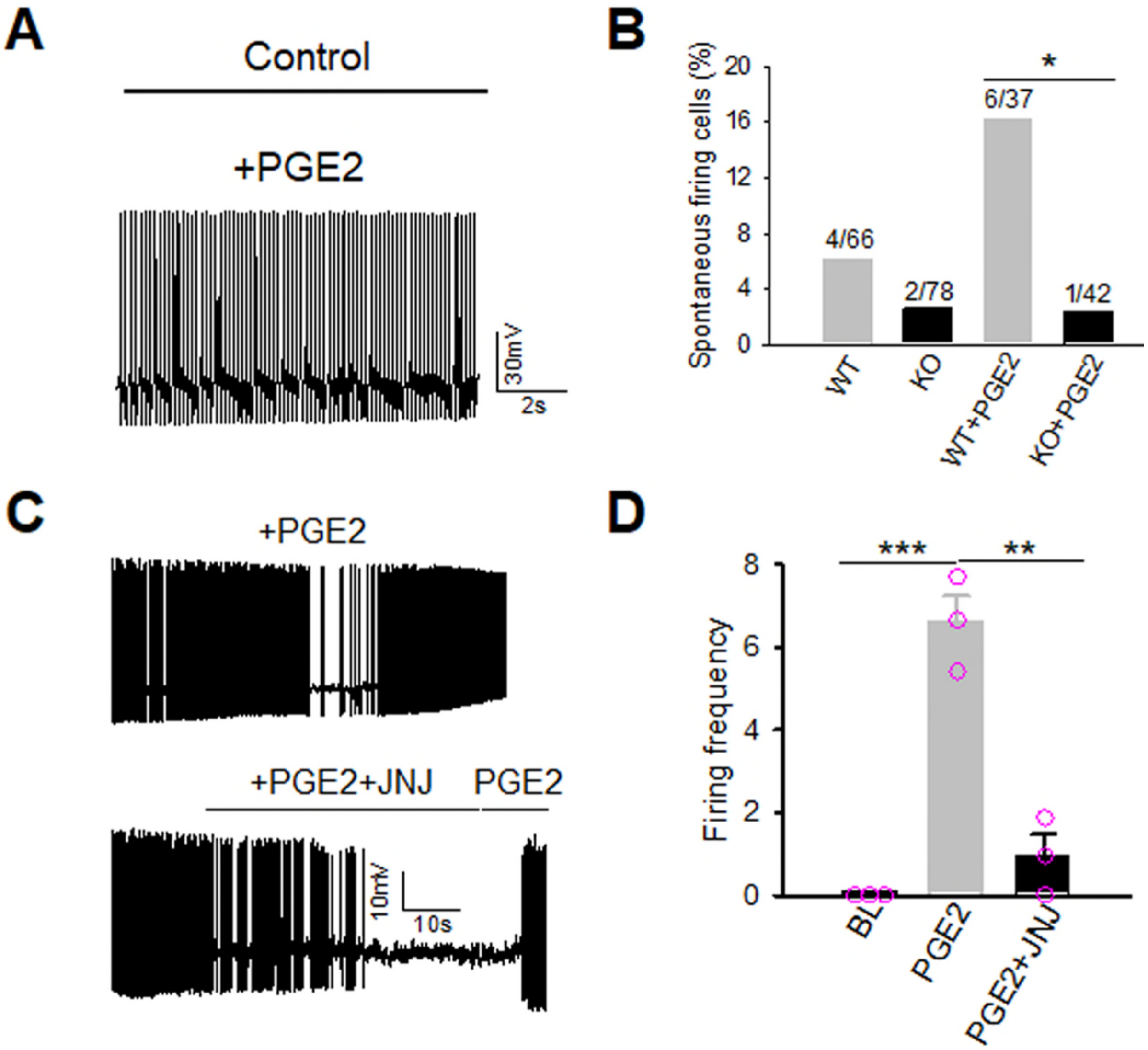


**Fig. S6: TRPM2 mediates spontaneous firing in DRG neurons evoked by PGE2. (A)** Example firing traces in a DRG neuron under control and after treatment with PGE2 (2μM, 3min). (**B**) Summary of the number of DRG neurons from WT and TRPM2-KO mice exhibiting spontaneous firing before and after treatment with PGE2 from experiments similar to those in (**A**). (**C**) Example firing of DRG neurons elicited by PGE2 (2μM, top panel) and in the presence of JNJ-28583113 (10μM) (bottom panel). After removal of JNJ-28583113, PGE2 elicited firing again. (**D**) Summary of firing frequency (spikes/s) in DRG neurons before and after treatment with PGE2 and JNJ28583113 from experiments similar to those in (**C**).

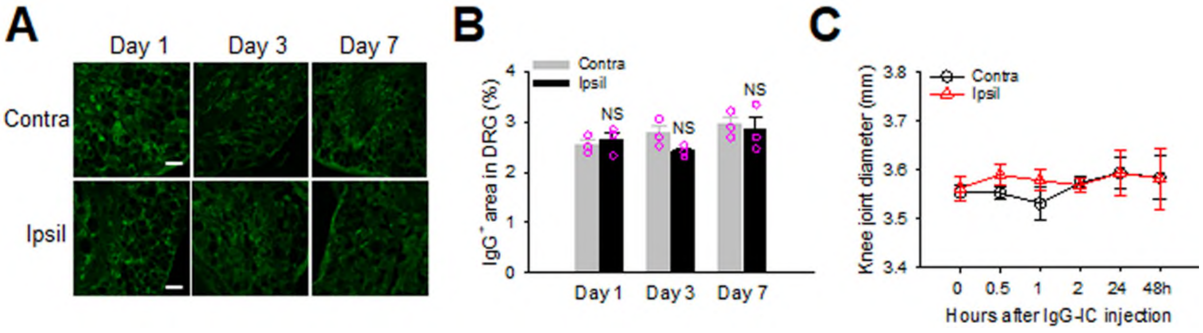


**Fig. S7: IgG is not increased in AIA model.** (**A**) Example fluorescence images of IgG immunoreactivity in the contralateral and ipsilateral DRG from AIA mice after different days of induction. Scale bars, 50μm. **(B)** Summary of IgG-positive areas in the DRG from similar experiments to those in (**A**). (**C)** Measurement of knee joint diameter at different time points after injection of IgG-IC (2μg). No significant joint inflammation is detectable. n=7.

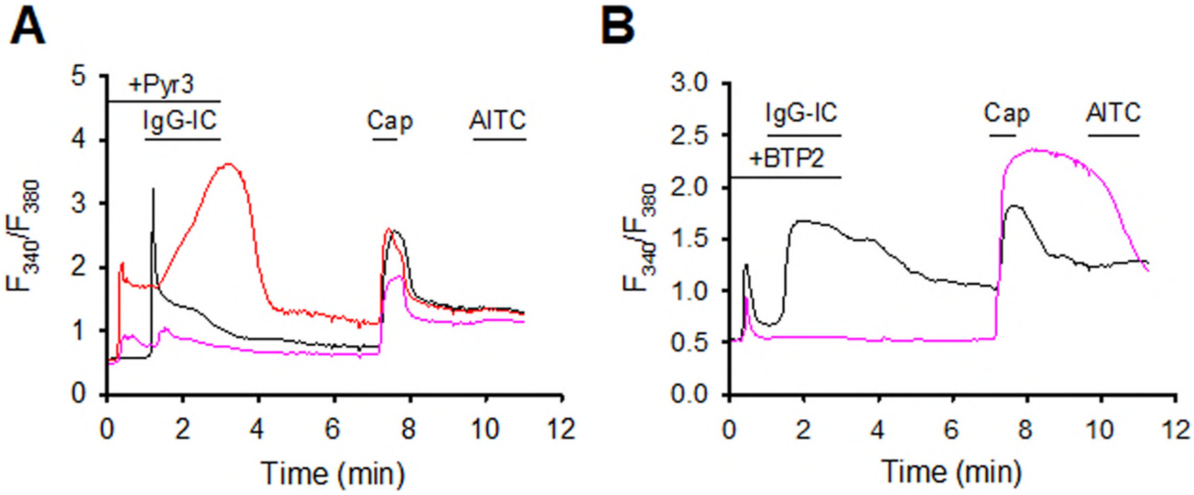


**Fig. S8: TRPC3 blockers elevate the basal $[Ca^{2+}]_i$ and enhance IgG-IC-induced $Ca^{2+}$ responses in DRG neurons. (A** and **B)** Example $Ca^{2+}$ responses in DRG neurons evoked by IgG-IC (1μg/ml) in the present of Pyr3 (10μM) (**A**) and BTP2 (10μM) (**B**). Both Pyr3 and BTP2 induced a basal $Ca^{2+}$ response and further enhanced IgG-IC-induced $Ca^{2+}$ responses.
